# The membrane-proximal domain of the periplasmic adapter protein plays a role in vetting substrates utilising channels 1 and 2 of RND efflux transporters

**DOI:** 10.1101/2021.10.05.463233

**Authors:** Ilyas Alav, Vassiliy N. Bavro, Jessica M. A. Blair

## Abstract

Active efflux by resistance-nodulation-division (RND) efflux pumps is a major contributor to antibiotic resistance in clinically relevant Gram-negative bacteria. Tripartite RND pumps, such as AcrAB-TolC of *Salmonella enterica* serovar Typhimurium, comprise of an inner membrane RND transporter, a periplasmic adaptor protein (PAP) and an outer membrane factor. Previously, we elucidated binding sites within the PAP AcrA (termed binding boxes) that were important for AcrB-transporter recognition. Here, we have refined the binding box model by identifying the most critical residues involved in PAP-RND binding and show that the corresponding RND-binding residues in the closely related PAP AcrE are also important for AcrB interactions. In addition, our analysis identified a membrane-proximal domain (MPD)-residue in AcrA (K366), that when mutated, differentially affects transport of substrates utilising different AcrB efflux-channels, namely channels 1 and 2, supporting a potential role for the PAP in sensing the substrate-occupied state of the proximal binding pocket (PBP) of the transporter and substrate vetting. Our model predicts that there is a close interplay between the MPD of the PAP and the RND transporter in the productive export of substrates utilising the PBP.

**Importance:** Antibiotic resistance greatly threatens our ability to treat infectious diseases. In Gram-negative bacteria, overexpression of tripartite efflux pumps, such as AcrAB-TolC, contributes to multidrug resistance because they export many different classes of antibiotics. The AcrAB-TolC pump is made up of three components: the periplasmic adaptor protein (PAP) AcrA, the RND-transporter AcrB, and the outer-membrane factor TolC. Here, we identified critical residues of AcrA that are important for its function with AcrB in *Salmonella enterica* serovar Typhimurium. Also, we show that AcrA shares these critical residues with AcrE, a closely related PAP, explaining their interoperability with AcrB. Importantly, we identified a residue in the membrane-proximal domain of AcrA that when mutated affected how different substrates access AcrB and impacted downstream efflux *via* TolC channel. Understanding the role that PAPs play in the assembly and function of tripartite RND pumps can guide novel ways to inhibit their function to combat antibiotic resistance.

## Introduction

Antibiotic resistance is one of the greatest global public health challenges and threatens our ability to effectively treat and prevent infectious diseases (1). In clinical isolates of Gram-negative bacteria, the Resistance-Nodulation-Division (RND) family of efflux pumps are frequently upregulated and associated with multidrug resistant phenotypes (2–6). Tripartite RND pumps span the double membrane of Gram-negative bacteria and consist of an inner membrane RND transporter, a periplasmic adaptor protein (PAP), and an outer membrane factor (OMF) (7, 8). The AcrAB-TolC pump is the principal RND efflux system in Enterobacteriaceae, including *Salmonella enterica*. It can export a wide range of structurally different compounds, including clinically relevant antibiotics such as β-lactams and fluoroquinolones (9, 10).

Gram-negative bacteria encode a wide repertoire of RND transporters, which typically pair with a single cognate PAP and an OMF to form tripartite pumps that have varied substrate specificities and physiological roles (10–16). The *S. enterica* genome encodes five RND pumps: AcrAB-TolC, AcrEF-TolC, AcrAD-TolC, MdtABC, and MdsABC (10). The AcrEF-TolC pump possesses a similar substrate profile to AcrAB-TolC, but its expression is silenced by H-NS under laboratory conditions (17). The PAPs comprise four domains: α-helical domain, lipoyl domain, β-barrel domain and the membrane-proximal domain (MPD) (18). The α-helical domain has a coiled-coil arrangement and appears to interact with the α-barrel domain of the OMF (18, 19). The lipoyl domain is involved in stabilising the self-assembly of the PAPs within the tripartite efflux pump. The β-barrel domain is flexibly linked to the MPD, and both domains appear to interact with the porter domain of the RND-transporter (20). The MPD of RND-associated PAPs is critical for the assembly and function of AcrAB-TolC in *E. coli* and *S. enterica* (21, 22). In the PAPs ZneB and CusB of the tripartite ZneCAB and CusABC RND efflux pumps, respectively, the MPD appears to play an important role in substrate acquisition and presentation to the metal pumping RND transporters (23, 24). Additionally, in the related ABC-transporter-associated PAP MacA, the MPD has been demonstrated to be involved in direct binding of possible pump substrates (25) and has been suggested to be involved in substrate vetting (18).

Previous studies have shown that AcrA and AcrE are interchangeable in *S. enterica* (24–26), whereas MdtA and MdsA can only function with their cognate RND-transporters (22). Previously, we showed that the regions of PAP-transporter contact are relatively compact and discrete. Based on homology models of the PAPs in *Salmonella*, we found these regions to be highly conserved between AcrA and AcrE, while differing significantly between divergent PAPs, such as MdtA and MdsA, providing a possible explanation of the observed interoperability of AcrA and AcrE. The 3D RND-interaction sites can be delineated into discrete linear sequences, which we have dubbed “binding boxes”, that map to the β-barrel domain (boxes 1–5) and the MPD (6–9). Disruption of a few key residues within the binding boxes 1 and 4-6 mapping to the exposed β-barrel loops and the MPD, abrogated transport, suggesting an important role for this region in AcrB-binding (22).

Here, we set out to further validate the “binding boxes” model of PAP-RND interaction by phenotypic profiling of site-directed mutants targeting the β-barrel and membrane-proximal domains. We specifically sought to describe the efflux profiles of substrates that have been suggested to utilise different AcrB-efflux channels (26–28).

## Results

### Refining the binding box model of PAP-RND interactions

Previously, using disruptive site-directed mutagenesis (SDM), we demonstrated that discrete stretches of residues, which we dubbed “binding boxes” based on their spacial proximity (19, 20, 29, 30), control PAP-RND complex formation and recognition of cognate PAP-RND pairs (22). Here, we set out to refine our binding box model by generating and testing the effects of more subtle and conservative mutations to identify the PAP-residues most critical for RND-binding (Fig. 1). Specifically, residues which were previously shown to be important for RND-binding (22) were mutated to residues with similar properties to produce conservative mutations, while residues, the previous mutation of which led to limited functional impact were subjected to more disruptive mutagenesis. Mutated versions of AcrA were expressed in the *Salmonella* SL1344 Δ4PAP strain, which lacks all four known RND-associated PAPs (AcrA, AcrE, MdtA and MdsA) (22). The effects of the mutations on efflux function were assessed using ethidium bromide efflux assays and antimicrobial susceptibility testing.

**Figure 1.**
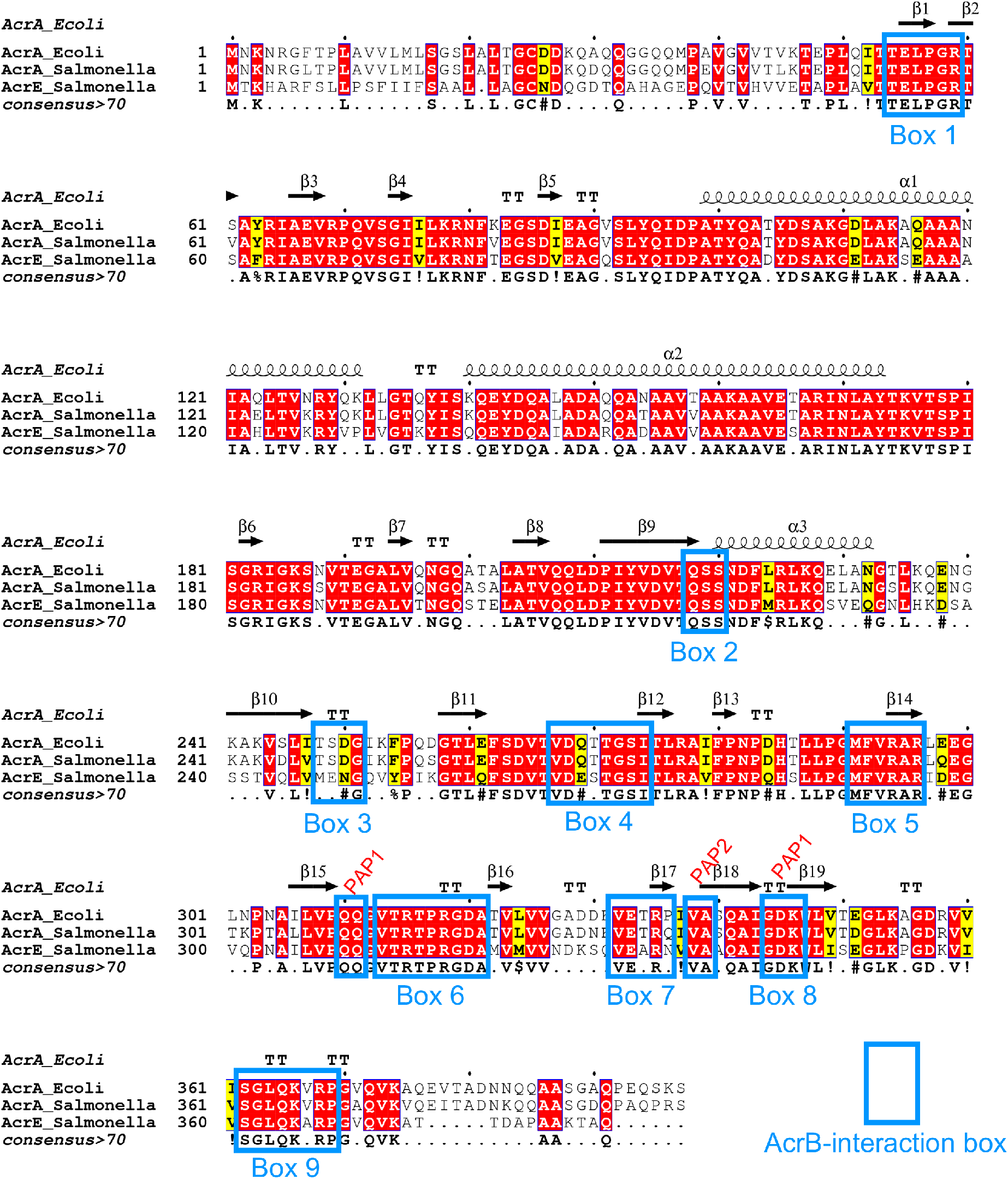
Multiple sequence alignment of *Salmonella* AcrA and AcrE combined with the mapping of the secondary structure derived from the experimentally defined structure of *E. coli* AcrA (PDB 5V5S, chain G) (20). Identical residues are coloured red and similar residues are coloured yellow. The PAP-binding boxes implicated in RND-binding (22) are numbered 1 to 9 and depicted using blue rectangles. Figure created using Espript 3.0 (34).

The PAPs reside in the periplasm and are embedded in the plasma membrane by a lipid anchor and/or a transmembrane helix and are composed of several well-defined domains (18, 31). From the plasma membrane outward, these are the membrane-proximal domain (containing boxes 6-9), the β-barrel domain (containing boxes 1-5), the lipoyl domain, and the α-hairpin domain (18). The G58F mutant mapping to box 1 was previously shown to impair efflux function (22). Here, the R59A mutant was produced to investigate whether other residues in proximity to box 1 had a role in RND-binding. The R59A mutant caused an intermediate impairment of efflux function, between that of the Δ4PAP and the WT complement strain, confirming the important role of box 1 (Fig. 2 and Table S2). Residues which were mutated as pairs in our previous study were separated to identify the most critical residue. The T270F-T271F mutant mapping to box 4 was separated into T270D and T271D for disruptive mutagenesis, and T270A and T271A mutants as more subtle mutations. The T271D mutant significantly impaired efflux, whilst the T270D mutant had no effect. Furthermore, the T271A mutation still caused a mild impairment of efflux function (Fig. 2 and Table S2), suggesting that T271 is a critical residue in efflux function. The G272P-S273P mutant mapping to box 4 previously impaired efflux function, therefore, a more conservative mutation was produced to determine the role of G272-S273. The G272A-S273A mutant had no effect on efflux activity, indicating that these residues can tolerate neutral mutations (Fig. 2 and Table S2). The F292G mutant mapping to box 5 was previously shown to affect both AcrB- and AcrD-binding (22, 32). Hence, the F292V mutant was produced to determine whether a more subtle change would still affect efflux function. Like the F292G phenotype, the F292V mutant also resulted in significantly abrogated efflux (Fig. 2 and Table S2), indicating that F292 is critical for efflux function. The Q310F mutant mapping to box 6 was previously shown to have no impact on efflux function, and the Q311F reported here, similarly did not influence efflux function detectably (Fig. 2 and Table S2). Therefore, both residues were simultaneously mutated into Q310P-Q311P, which resulted in significant impairment of efflux function, indicating that double glutamine residues may provide some functional redundancy and that the presence of a glutamine residue may be critical in this position (Fig. 2 and Table S2).

**Figure 2.**
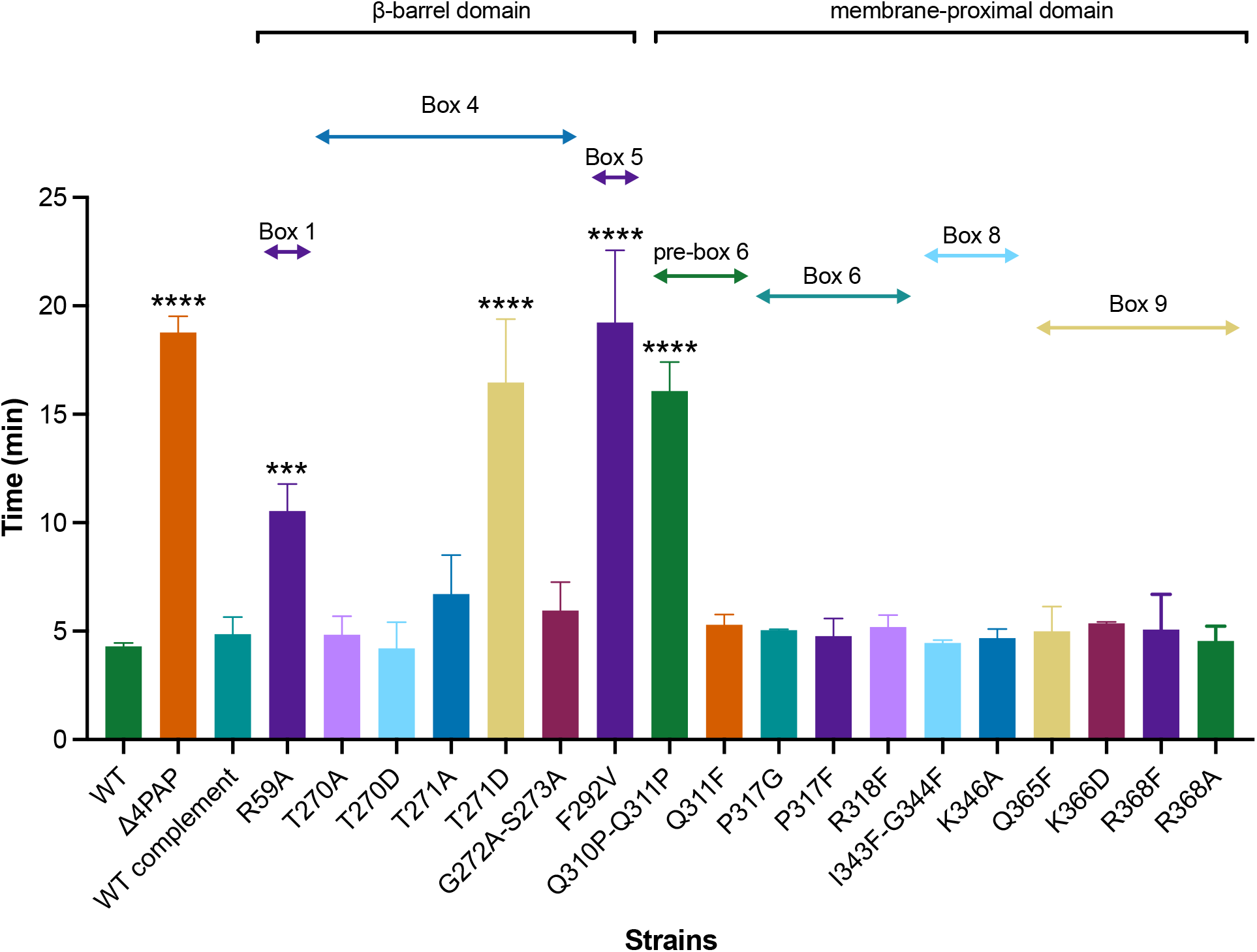
Efflux of ethidium bromide by the Δ4PAP strain complemented with mutated versions of AcrA. Data presented are the mean of three biological replicates and are shown as the time taken for the fluorescence to decrease by 25% +/− SD. Bacteria were treated with ethidium bromide and the proton-motive force dissipator CCCP for 1 hour and then re-energised with glucose. Annotation above indicates the mapping of each mutation to its binding box, as well as the domain mapping of respective boxes. Data were analysed by one-way ANOVA and compared to the WT complement using Dunnett’s test. Significantly different strains are denoted with *** (*P* ≤ 0.001) or **** (*P* ≤ 0.0001).

Mutations mapping to box 6, including R315F and R318A, were previously shown to have no impact on efflux function. Therefore, several novel mutations mapping to box 6 were produced (P317G, P317F and R318F), which did not have any impact on efflux function (Fig. 2 and Table S2). Novel mutations mapping to box 8 (I344F-G344F and K346A) also had no observable influence on efflux function (Fig. 2 and Table S2). G363 in box 9 was shown by us and other studies to be critical for efflux function (22, 32, 33). Further mutations in box 9 were produced to investigate whether other residues also play a role in RND-binding. However, Q365F, K366D, R368F and R368A had no effect on efflux function, suggesting that only G363 is critical for efflux function (Fig. 2 and Table S2). Western blotting verified that the observed effects of the mutants with impaired efflux were not due to changes in protein expression levels or stability (Fig. S1A).

### AcrE and AcrA share conserved binding boxes that are responsible for their interoperability relative to AcrB

The binding boxes between AcrA and AcrE were previously shown to be highly conserved in *Salmonella* Typhimurium (Fig. 1), potentially explaining the observed interchangeability between the two PAPs (22). To validate their functional role in AcrE, SDM was used to mutate the residues, corresponding to the most critical binding box residues previously identified in AcrA (22) - namely G57, R58, T270, F291, R293 and G362 (Fig. 1). The effect of the mutations was assessed by ethidium bromide accumulation assays and antimicrobial susceptibility testing in the Δ4PAP Δ*acrF* strain (22). This strain lacks all four RND-associated PAPs and the cognate RND-transporter AcrF, thereby allowing the impact of AcrE mutations on AcrB-binding to be determined. All mutations corresponding to the critical residues of AcrA (AcrE G57F, R58A, T270D, F291G, R293F and G362F) also had a significant effect on efflux function and antimicrobial susceptibility (Fig. 3 and Table S3). Consistent with this, the mutation of phenotypically neutral residues in AcrA, corresponding to the AcrE T216F, K365D and R367D respectively, also had no impact on efflux function (Fig. 3 and Table S3). The observed effects of the mutations tested stemmed from their impact on the function of the protein and were not due to changes in expression levels or stability of the variant alleles, as validated by Western blotting, with a possible exception of G57F (Fig. S1B).

**Figure 3.**
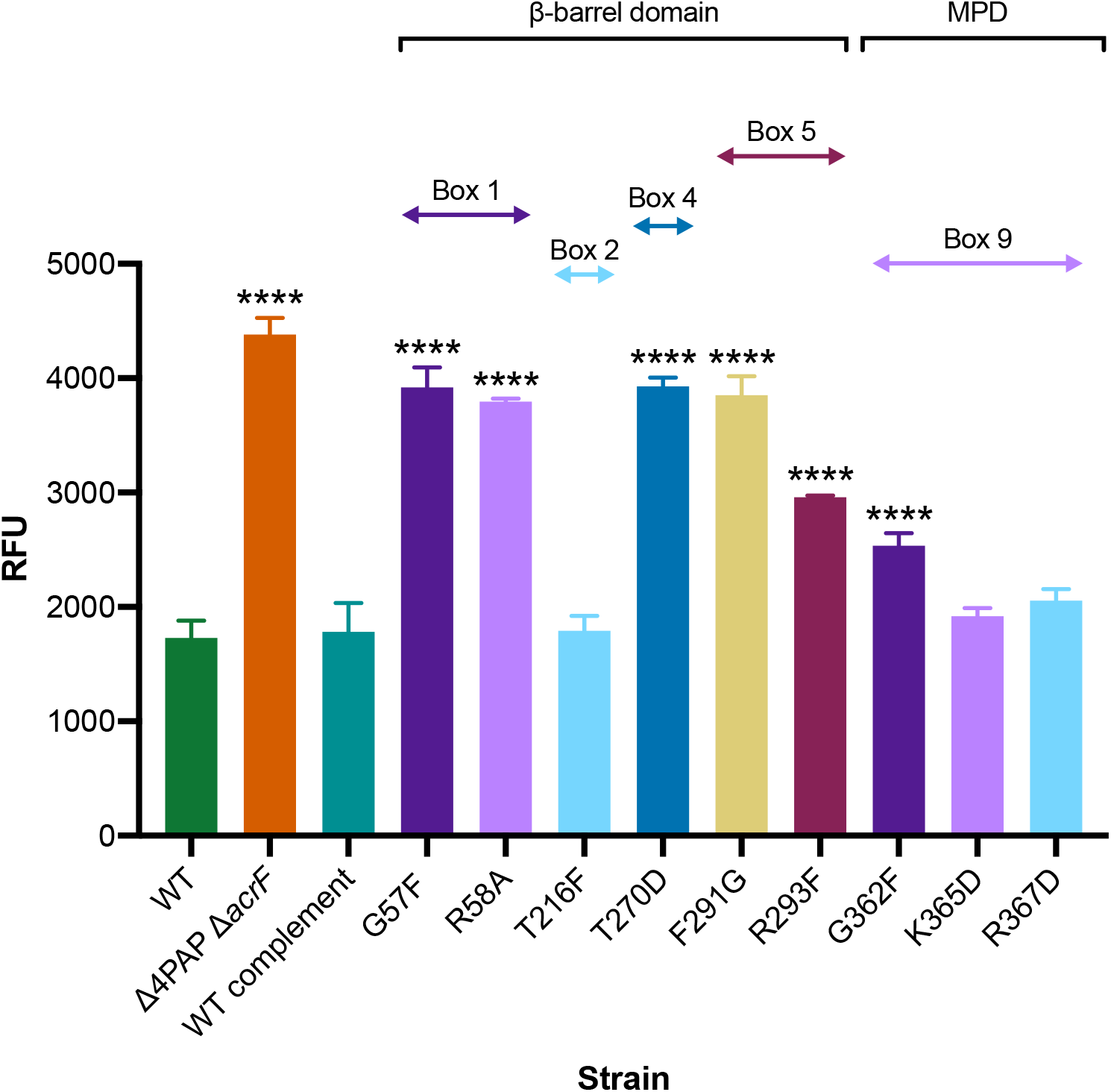
Accumulation of ethidium bromide in Δ4PAP ΔacrF strain complemented with mutated versions of AcrE. Data represented are the mean of three biological replicates showing maximum RFU values after 30 min of ethidium bromide exposure +/− SD. Annotation above indicates the mapping of each mutation to its binding box, as well as the domain mapping of respective boxes. Data were analysed by one-way ANOVA and compared to the WT complement strain using Dunnett’s test. Significantly different strains are indicated with **** (*P* ≤ 0.0001). MPD, membrane-proximal domain.

The above results confirm the conservation of function of the binding boxes between AcrA and AcrE, which explains their interoperability in conjunction with AcrB.

### Potential role for the membrane-proximal domain of AcrA in vetting substrate access to channel 1 and channel 2

The refinement of the binding box model of PAP-RND interaction led to the discovery of an AcrA mutant with a peculiar phenotype. The K366D AcrA mutant mapping to box 9 (Fig. 1), did not alter ethidium bromide efflux or susceptibility (Fig. 2), but showed a distinct antimicrobial susceptibility profile to other antimicrobials tested (Table S2). Notably, the K366D mutation in AcrA conferred differential effects depending on the physicochemical properties of the compounds tested. The K366D mutant displayed greater than two-fold reduction in MIC values to high-molecular-mass drugs (HMMDs), such as doxorubicin, erythromycin, fusidic acid and novobiocin (*M >* 500 g/mol) and low-molecular-mass drugs (LMMDs), such as chloramphenicol, clindamycin, linezolid, and minocycline (*M <* 500 g/mol), compared to WT AcrA complement. However, the K366D mutant had no impact on the MIC values for planar aromatic cations (PACs), including acriflavine, berberine, benzalkonium chloride, crystal violet, ethidium bromide, methylene blue and rhodamine 6G (Table 1).

**Table 1.**
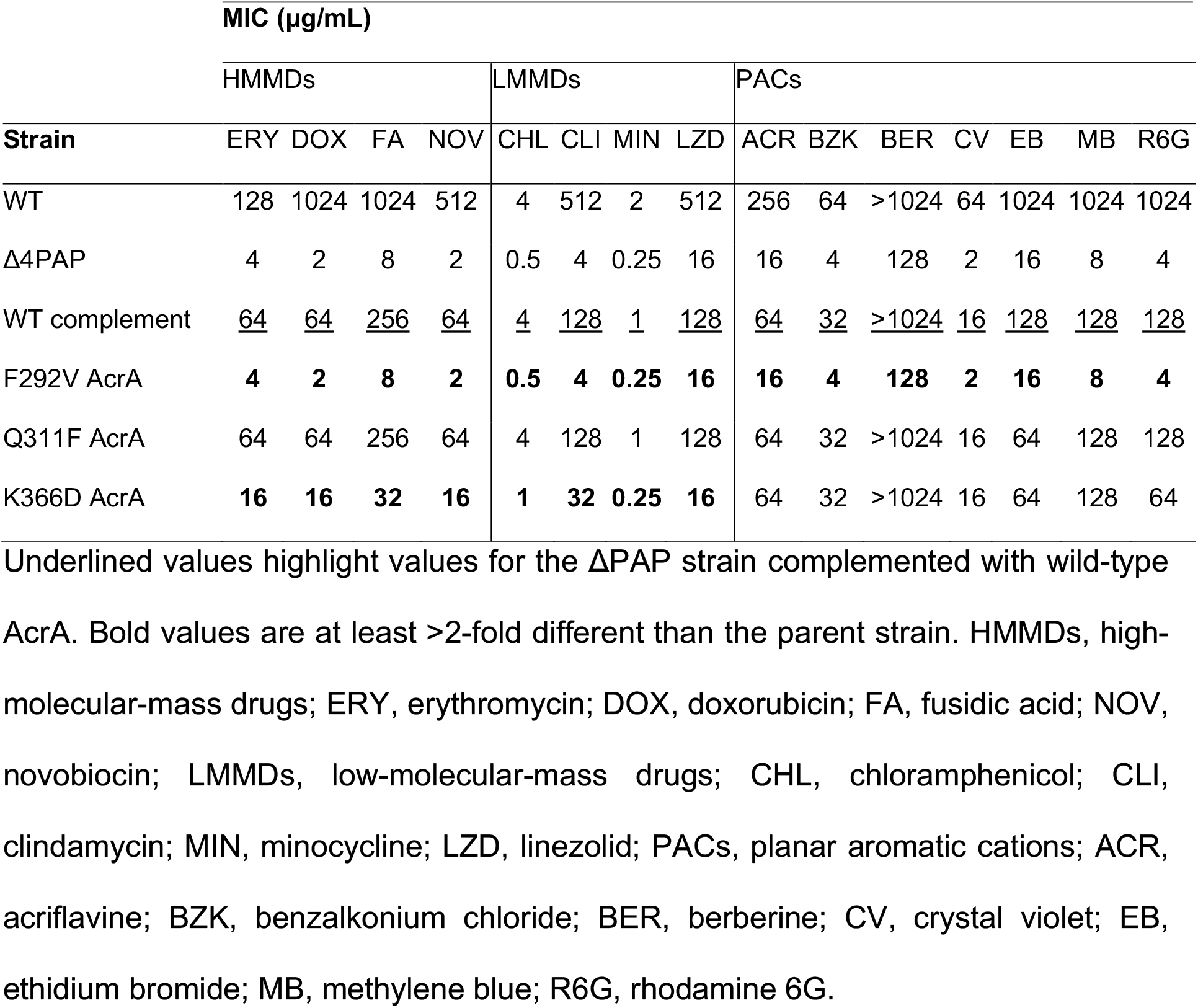
Antimicrobial susceptibility of Δ4PAP strain complemented with F292V, Q311F or K366D AcrA to drugs with different physicochemical characteristics.

| Strain | MIC (μg/mL) | | | | | | | | | | | | | | |
| --- | --- | --- | --- | --- | --- | --- | --- | --- | --- | --- | --- | --- | --- | --- | --- |
|  | HMMDs |  |  |  | LMMDs |  |  |  | PACs |  |  |  |  |  |  |
|  | ERY | DOX | FA | NOV | CHL | CLI | MIN | LZD | ACR | BZK | BER | CV | EB | MB | R6G |
| WT | 128 | 1024 | 1024 | 512 | 4 | 512 | 2 | 512 | 256 | 64 | >1024 | 64 | 1024 | 1024 | 1024 |
| Δ4PAP | 4 | 2 | 8 | 2 | 0.5 | 4 | 0.25 | 16 | 16 | 4 | 128 | 2 | 16 | 8 | 4 |
| WT complement | <u>64</u> | <u>64</u> | <u>256</u> | <u>64</u> | <u>4</u> | <u>128</u> | <u>1</u> | <u>128</u> | <u>64</u> | <u>32</u> | <u>>1024</u> | <u>16</u> | <u>128</u> | <u>128</u> | <u>128</u> |
| F292V AcrA | **4** | **2** | **8** | **2** | **0.5** | **4** | **0.25** | **16** | **16** | **4** | **128** | **2** | **16** | **8** | **4** |
| Q311F AcrA | 64 | 64 | 256 | 64 | 4 | 128 | 1 | 128 | 64 | 32 | >1024 | 16 | 64 | 128 | 128 |
| K366D AcrA | **16** | **16** | **32** | **16** | **1** | **32** | **0.25** | **16** | 64 | 32 | >1024 | 16 | 64 | 128 | 64 |

Underlined values highlight values for the $\Delta$ PAP strain complemented with wild-type AcrA. Bold values are at least >2-fold different than the parent strain. HMMDs, high-molecular-mass drugs; ERY, erythromycin; DOX, doxorubicin; FA, fusidic acid; NOV, novobiocin; LMMDs, low-molecular-mass drugs; CHL, chloramphenicol; CLI, clindamycin; MIN, minocycline; LZD, linezolid; PACs, planar aromatic cations; ACR, acriflavine; BZK, benzalkonium chloride; BER, berberine; CV, crystal violet; EB, ethidium bromide; MB, methylene blue; R6G, rhodamine 6G.

Previous studies have associated molecular weight of the substrate-drugs with the preferred channel access and binding pocket validation by the RND-transporter (26–28, 35). The RND-transporter AcrB has multiple substrate entry channels identified, which are used by drugs depending on their physicochemical properties (26, 27, 36) (Fig 4). LMMDs have been proposed to preferentially enter through channel 1 (CH1), whilst HMMDs are thought to enter AcrB through channel 2 (CH2) (36–38). The cleft entrance of CH2 has been previously suggested to be impacted by the membrane-proximal domain (MPD) of AcrA (26). This is seen in cryo-EM structures of the AcrAB-TolC complex, which show that AcrA interacts with the PC1 and PC2 subdomains of AcrB (19, 39). PACs on the other hand, are preferentially taken up through channel 3 (CH3), which starts from the vestibule formed by the central cavity of the three AcrB protomers and leads directly to the deep binding pocket (DBP) (26). The entrance of the recently proposed channel 4 (CH4) is in the groove formed by TM1 and TM2 and leads to the DBP (40). The location of CH3 and CH4 within AcrB suggests that the MPD of AcrA should not have a direct steric impact on the drug access (Fig. 4).

**Figure 4.**
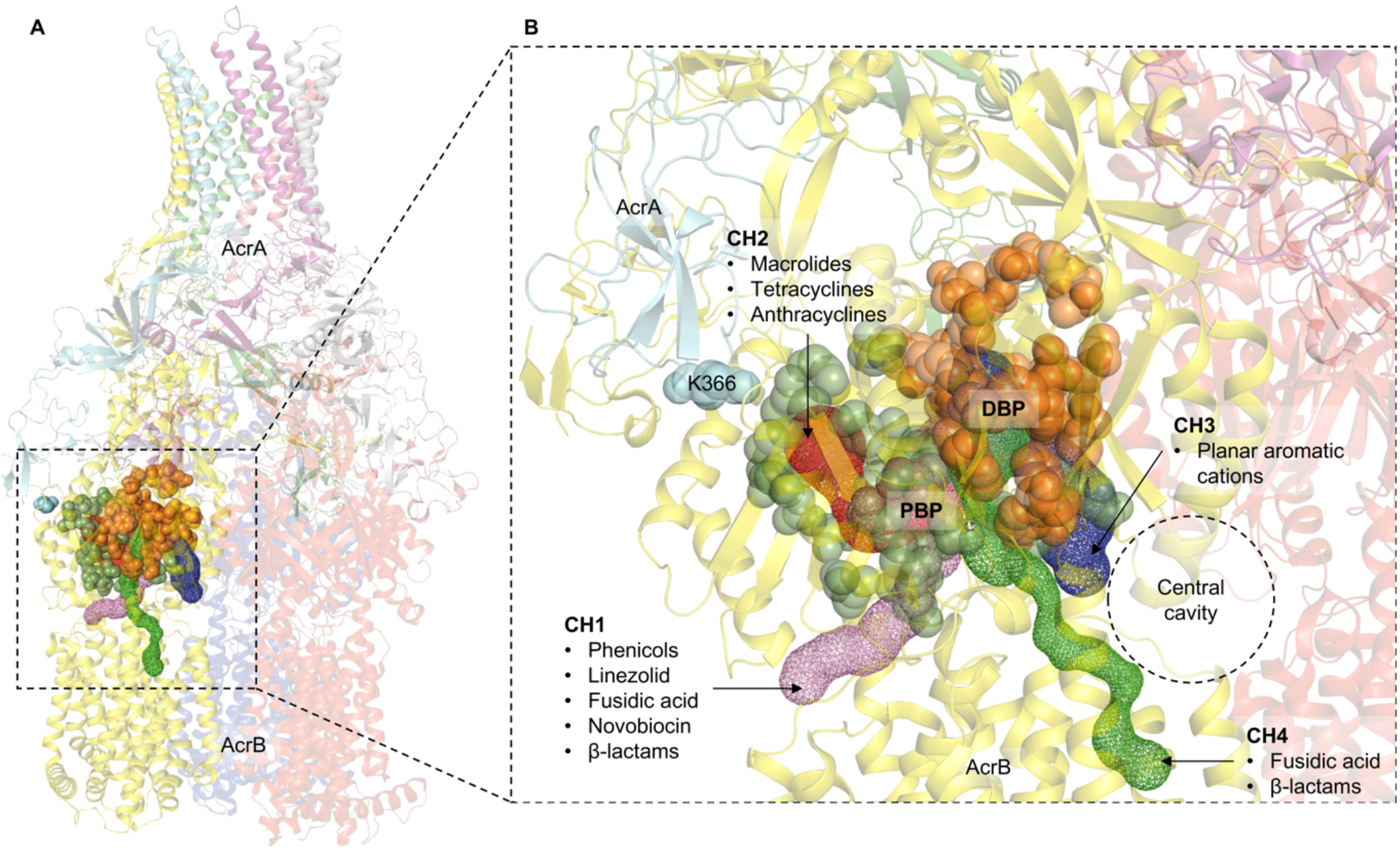
**A.** The crystal structure of the trimeric AcrB transporter and the hexameric AcrA assembly (TolC not shown for clarity). The different substrate entry pathways are shown as coloured channels, and the binding pockets are indicated by coloured spheres. **B.** Zoomed-in view of the substrate channels and the binding pockets relative to K366 of AcrA. The green and orange spheres correspond to the space-fill representation of the residues lining the proximal binding pocket (PBP) and the deep binding pocket (DBP), respectively. K366 is in the membrane-proximal domain of AcrA and impacts the residues lining the PBP and the entrance of channel 2 (CH2). Channel 1 (CH1) also feeds into the PBP, so is likely to be impacted by changes in K366. Channel 3 (CH3) starts from the central cavity and leads to the DBP. Similarly, channel 4 (CH4) starts from the groove formed by TM1/TM2 and leads to the DBP. Therefore, CH3 and CH4 are unlikely to be directly impacted by K366 substitutions.

Therefore, to understand how the K366D AcrA mutation affects the susceptibility of the Δ4PAP strain to HMMDs and LMMDs, but not to PACs, we designed more specific antimicrobial sensitivity screens to be able to better differentiate the usage of the access channels by respective substrates and the impact of the K366D mutation on specific channels.

**Figure 5.**
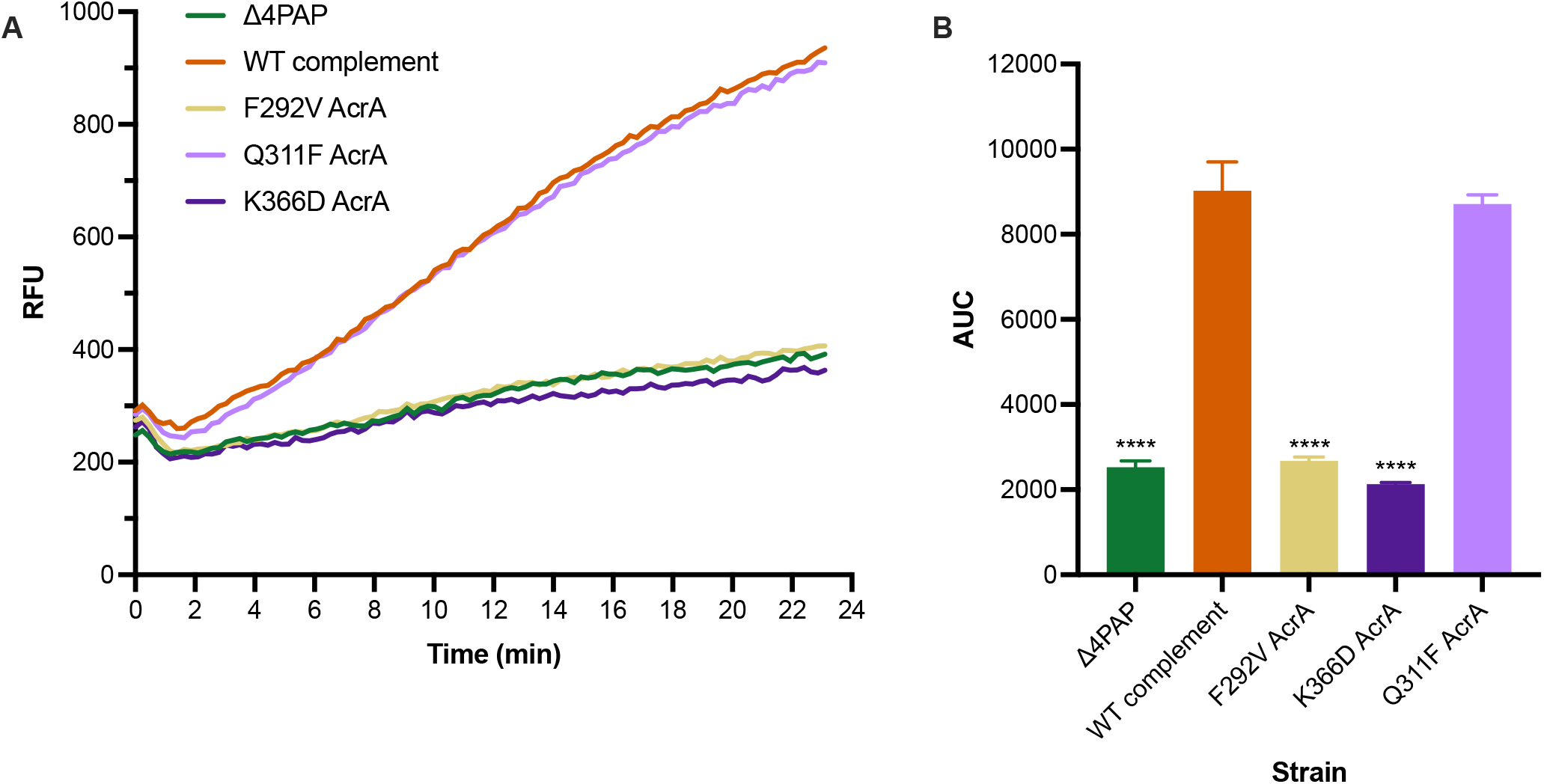
A) Efflux of doxorubicin over time in Δ4PAP strain complemented with mutated versions of AcrA. Bacteria were treated with doxorubicin and the proton-motive force dissipater CCCP for one hour and then re-energised with glucose. Efflux was monitored by increasing RFU due to extracellular doxorubicin. Data presented are the mean of three biological replicates. **B) Area under curve (AUC) analysis for doxorubicin efflux over time.** Data shown are the mean AUC of the three biological replicates shown in panel A. Data were analysed by one-way ANOVA and compared to the WT complement using Dunnett’s test. Strains with a significantly different AUC are indicated with **** (*P* ≤ 0.0001).

To further clarify the impact of the K366D mutation on specific channels, we monitored growth in the presence of efflux-substrates at 0.25×MIC for K366D AcrA, using the efflux-impaired F292V and phenotypically neutral Q311F as controls. The growth kinetics data showed that, compared to the WT complement, the K366D AcrA mutant grew poorly or not at all in the presence of CH2 substrates doxorubicin, erythromycin, minocycline, and clindamycin (Fig. 6). The K366D AcrA mutant also displayed growth defects when grown in CH1 substrates, including novobiocin, fusidic acid and chloramphenicol (Fig. 6). Notably, the K366D AcrA mutant did not have any observable growth defects when grown in CH3 substrates, such as ethidium bromide and rhodamine 6G (Fig. 6). This data shows disproportionate impact of the K366D on substrates utilising CH1 and CH2 (27, 28), while substrates documented to utilise CH3, such as ethidium bromide and rhodamine 6G (26) appear relatively unaffected.

**Figure 6.**
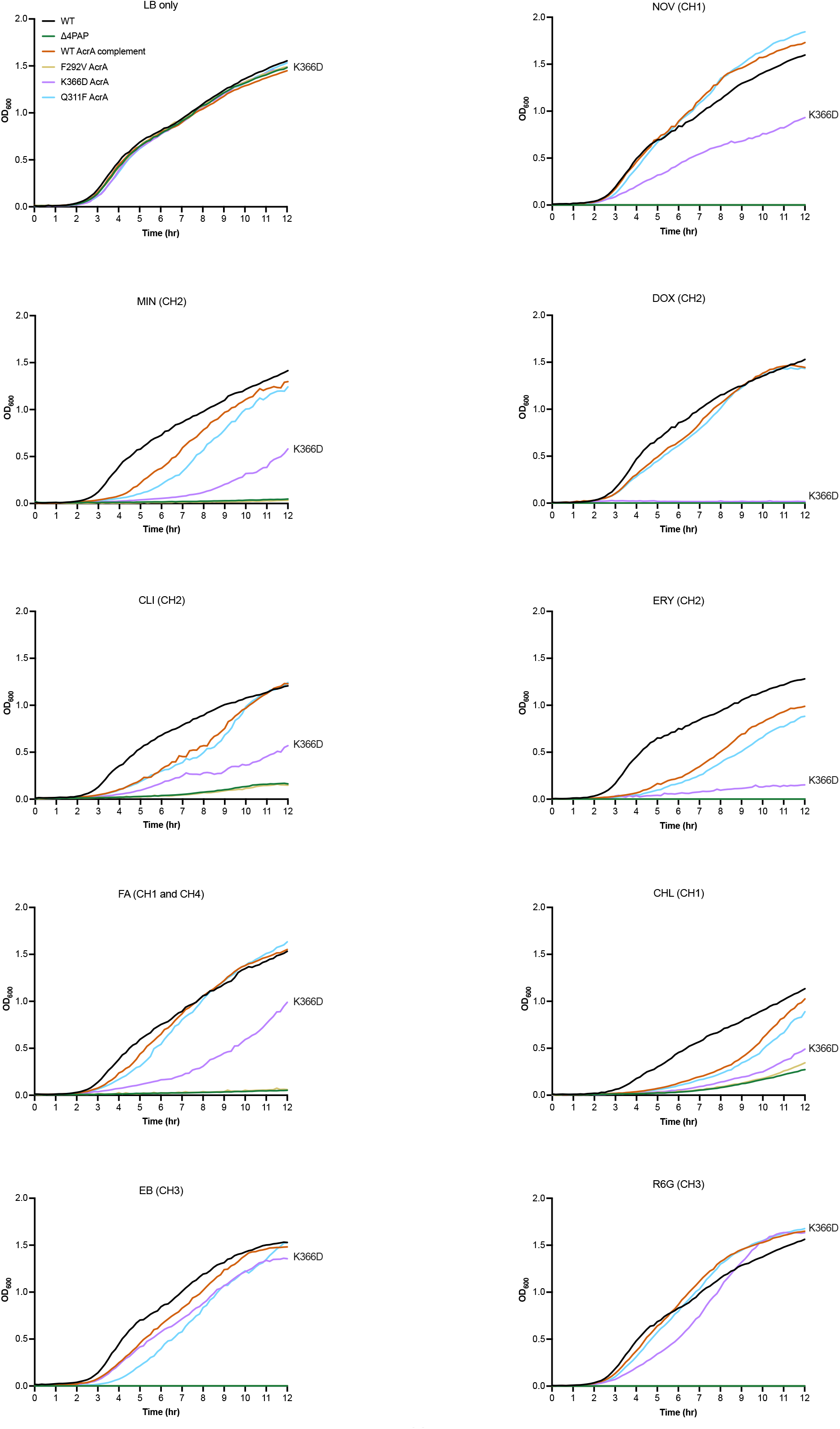
Growth kinetics of Δ4PAP strain complemented with mutated versions of AcrA. Abbreviations and concentrations of drugs used: CHL, 0.5 μg/mL chloramphenicol; CLI, 8 μg/mL clindamycin; DOX, 4 μg/mL doxorubicin; EB, 16 μg/mL ethidium bromide; ERY, 4 μg/mL erythromycin; FA, 8 μg/mL fusidic acid; MIN, 0.25 μg/mL minocycline; NOV, 4 μg/mL novobiocin; R6G, 16 μg/mL rhodamine 6G. Brackets indicate the preferred channel utilised by the substrate: CH1, channel 1; CH2, channel 2; CH3, channel 3; CH4, channel 4. Data shown are the mean OD_600_ values of three biological replicates. Concentrations of drugs are 0.25×MIC of K366D AcrA.

### AcrB mutant with impacted channel 3 function supports the role of K366 in ensuring productive efflux of channel 1 and channel 2 substrates

The above data, combined with the location of the K366D mutation, strongly suggests that it may affect substrates entering through CH1 and CH2, but not CH3. To further increase assay sensitivity and avoid interference from substrates that use CH1-3 promiscuously, such as ethidium bromide, we used an AcrB mutant (A33W, T37W, N298W AcrB) with impacted CH3 function (26), which allows for better separation of efflux signal arising from CH1 and CH2. An AcrB CH3 mutant (A33W T37W N298W AcrB), which was previously shown to impact the export of PACs (26). The AcrB CH3 mutant displayed an intermediate level of efflux impairment between that of Δ4PAP Δ*acrB* and the WT *acrAB* complement strain (Fig 7). Furthermore, the AcrB CH3 mutant displayed increased susceptibility to PACs (Table S4) and impaired growth in the presence of 32 μg/mL ethidium bromide (Fig. S2). Importantly, when present in combination with the AcrB CH3 mutant, the K366D mutation abrogated ethidium bromide efflux even further, to a level comparable to that of the Δ4PAP Δ*acrB* strain (Fig. 7). Antimicrobial susceptibility testing also showed that the K366D AcrA with the AcrB CH3 mutation displayed increased susceptibility to substrates compared to the K366D AcrA or AcrB CH3 mutants alone (Table S4).

**Figure 7.**
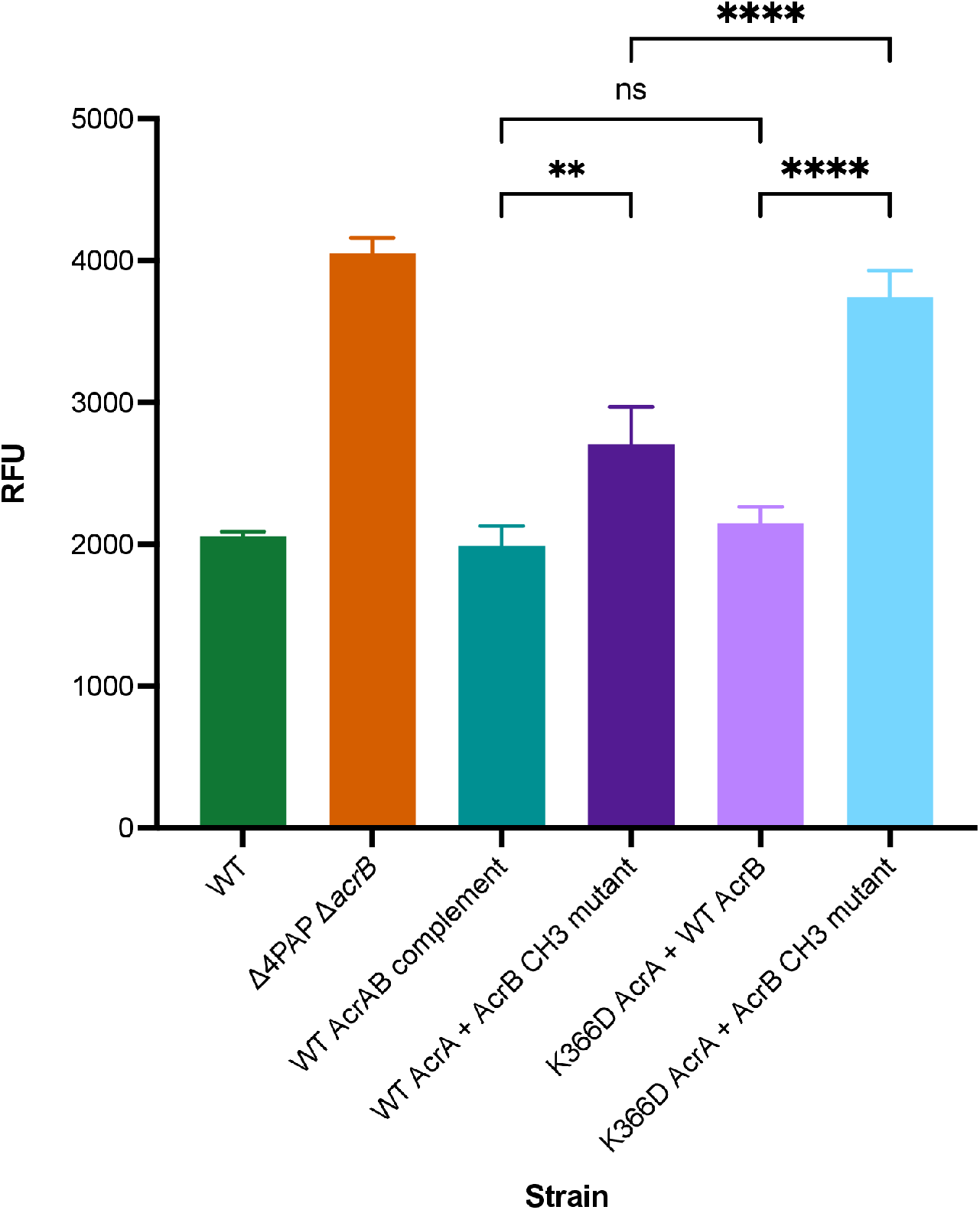
Accumulation of ethidium bromide in Δ4PAP Δ*acrB* strain complemented with K366D AcrA and the AcrB CH3 mutant (A33W T37W N298W AcrB). Data shown are the mean of three biological replicates showing maximum RFU values after 30 minutes of ethidium bromide exposure. Data were analysed by one-way ANOVA and corrected for multiple comparisons using Tukey’s test. Significantly different strains are indicated with ** (*P* ≤ 0.01) or **** (*P* ≤ 0.0001). ns, not significant.

Next, the ability of the double mutant (K366D AcrA + AcrB CH3 mutant) to export doxorubicin was measured. The WT AcrA combined with the AcrB CH3 mutations showed a similar level of doxorubicin efflux as the WT AcrAB complement, whilst the K366D AcrA mutation with WT AcrB strain displayed impaired doxorubicin efflux. The double mutant showed complete impairment of doxorubicin efflux, like that of the Δ4PAP Δ*acrB* strain (Fig. 8). The doxorubicin efflux assay results were further validated by growing the K366D AcrA and the AcrB CH3 mutant strains in the presence of doxorubicin. The double mutant failed to grow in the presence of 2 μg/mL doxorubicin like the Δ4PAP Δ*acrB* strain. The K366D AcrA mutant with WT AcrB displayed impaired growth in the presence of 8 μg/mL doxorubicin, whilst the AcrB CH3 mutant with WT AcrA had no observable growth defect (Fig. S3). The concentration dependent effect of doxorubicin growth inhibition is consistent with the blockage of CH2, and gradual saturation of CH1, the function of which is partially impacted by K366D mutation. Western blotting verified that the phenotypic effects of the AcrB CH3 mutation were not due to changes in protein expression or stability (Fig. S1C). In summary, the AcrB CH3 disruption has a clearly pronounced additive effect compared to K366D acting on its own, consistent with the role of the MPD of the PAP in the control of CH1 and CH2 substrates. These data further support the essential role that K366, and the MPD in general play in the transport of CH1 and CH2 substrates.

**Figure 8.**
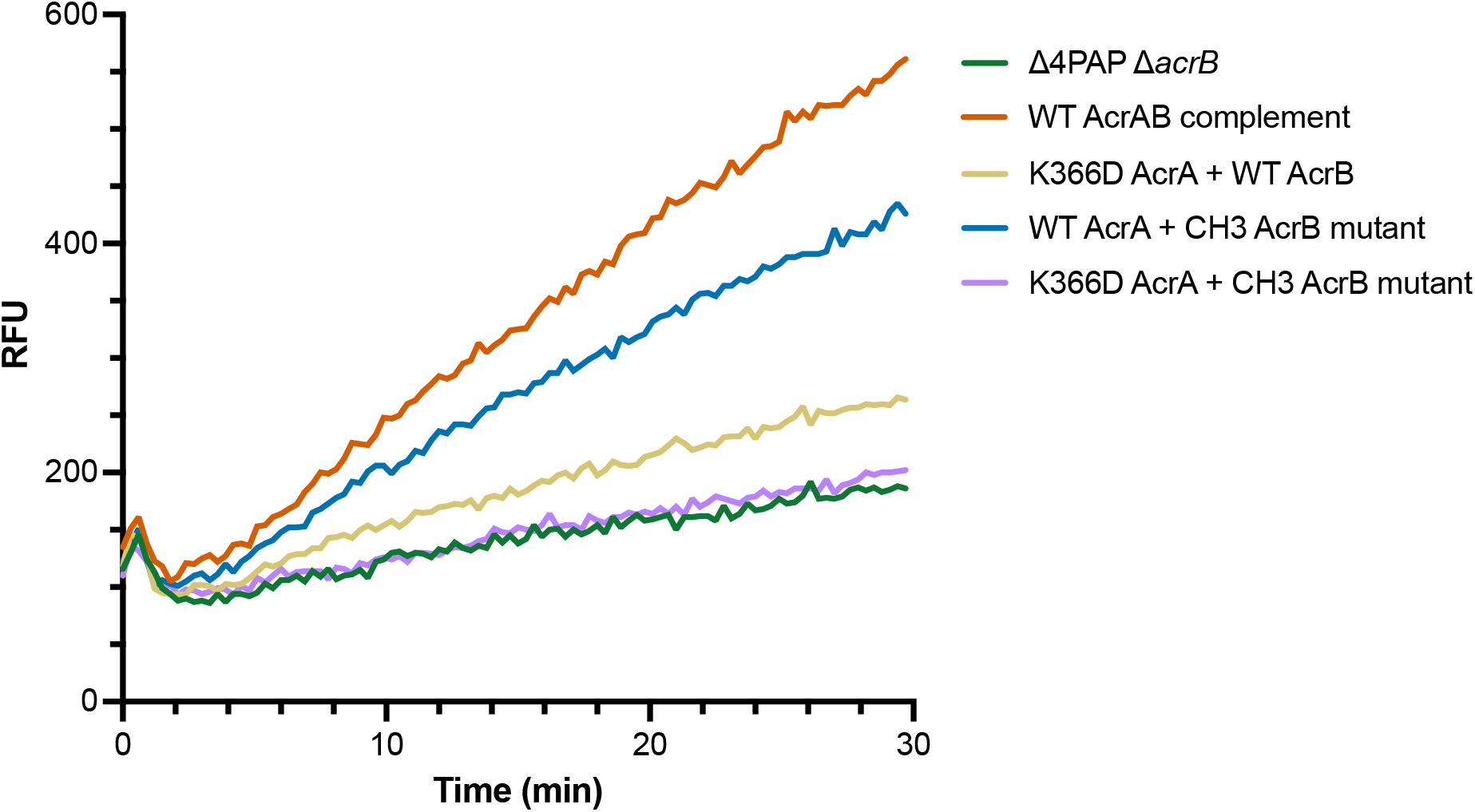
The effect of the K366D AcrA and the AcrB channel 3 (CH3) mutation on doxorubicin export. Efflux of doxorubicin over time in Δ4PAP Δ*acrB* strain complemented with K366D AcrA and the AcrB channel 3 (CH3) mutant. Bacteria were treated with doxorubicin and the proton-motive force dissipater CCCP for 1 hour and then re-energised with glucose. Efflux was monitored by increasing RFU due to extracellular doxorubicin. Data presented are the mean of three biological replicates. AcrB CH3 mutant refers to A33W T37W N298W AcrB.

### K366D also impacts channel 1 substrate transport

Like chloramphenicol, linezolid also uses CH1 (28) and consistent with the interpretation that K366D impacts efflux through this channel, we observed a similar result for linezolid, with clearly pronounced concentration-dependent effect, which is most pronounced at 8 μg/mL and above (Fig. S4). This data suggests that K366 is also somehow involved in either active substrate vetting or surveillance of the substrate-bound state of the transporter, as discussed in detail in the section below.

## Discussion

In this study, we set out to refine the previously reported “binding box” model of PAP-RND interaction (22) by generating and characterising additional subtle and more conservative PAP mutations. Consistent with the model’s prediction, we report that the AcrE residues that correspond to the previously identified critical residues in AcrA conserve their functional significance, as evidenced by targeted mutagenesis. Furthermore, we have been able to refine the effect of the previous, rather blunt mutations by separating previously reported double mutant AcrA T270F-T271F into AcrA T270A, T270D, T271A and T271D, which enabled us to narrow down the T271 as a crucial residue for efflux function. Likewise, the function of F292 also seems to be critical for efflux function, with even subtle mutations (AcrA F292V) resulting in complete destabilisation of the AcrAB-TolC assembly. The R59A mutant, which is in proximity to G58, resulted in intermediate impairment of efflux function, validating the previously reported role for box 1, and the PAP β-barrel domain in functional tripartite pump assembly.

During the refinement of the binding box 9, which belongs to the membrane-proximal domain of the PAP, we identified an AcrA mutant (K366D) with a peculiar phenotype. The K366D mutant had no impact on ethidium bromide efflux or susceptibility to PACs. However, it had significantly increased susceptibility to HMMDs and LMMDs compared to WT complement. Structural analysis of the available RND tripartite assemblies (20, 29, 30) indicated that K366 is in proximity to the proposed entry of the CH2 (Fig. 4). We hypothesised that if this were indeed the case, as substrates and drugs exhibit clear channel access preferences, K366D will disproportionately affect substrates using CH1 and CH2, but not CH3 or CH4. Consistent with this prediction, we observed that the K366D mutant exported doxorubicin, a CH2-substrate, very poorly. Furthermore, the K366D mutant displayed growth defects in the presence of several additional CH1-substrates (chloramphenicol, fusidic acid, and linezolid), as well as CH2-substrates (doxorubicin, erythromycin, and minocycline), but not CH3-substrates (ethidium bromide and rhodamine 6G).

Notably, the disruption of CH3 in AcrB had a clearly pronounced additive effect on the AcrA K366D mutant acting in a WT AcrB background, consistent with the role of the PAP MPD in the control of CH1 and CH2 access to the respective substrates. Intriguingly, in addition to the straightforward effect of K366D on the CH2 entry, the linezolid data presented here suggests that there is also a measurable impact on the CH1-substrates. This necessarily requires some level of allosteric communication, because K366 is too far from the proposed entrance of CH1 (26, 27, 41).

At this stage, the available data doesn’t provide a definitive answer as to how the MPD of the PAP may impact on the apparent substrate preference and selection. However, the analysis of the available PAP-RND complex structures (20, 29, 30) (Fig. 4), provides hints to the possible mechanism of the K366 action. One straightforward possibility, based on the location of K366 near the suggested entry of CH2 (26), and the flexibility of its side-chain is that it may affect CH2 substrate access and kinetics by playing the role of a “cap” on the tunnel entrance, possibly sensing, and even partially coordinating the incoming substrate. However, the effect of K366D also extends to CH1-substrates, such as linezolid and chloramphenicol, while K366 is located too far away from the suggested CH1 entry points to be directly involved in any active substrate vetting. Thus, it is tempting to suggest that the K366, and the MPD as a whole may be involved in a more generalised sensing of the substrate occupied state of the PBP (which is the convergence of CH1 and CH2) (27, 41), and/or the potential propagating of the “substrate-occupied” signal upwards *via* conformational change in the PAP leading to TolC engagement and channel opening as previously suggested (8). Strikingly, this interpretation is directly supported by the very recent *in situ* cryo-electron tomography structure of the assembled AcrAB-TolC pump (42), which displays strong and differential association of the MPDs of PAP protomers I and II with the PC1 and PN1/PC2 domains of AcrB respectively, with the latter in particular suggested to be associated with sensing the MBX3132 drug-occupied state of the transporter, and providing a conformational signal to TolC, affecting its channel gating. Additionally, for the first time, the *in-situ* structure also unambiguously identifies the location of the C-terminal helices and the membrane-associated N-terminal tails of AcrA (42), that appear to occupy a crevasse on the AcrB-surface that may also plausibly account for CH1 effects reported here, possibly providing additional sensory/allosteric input. Such sensory input may be allosterically conveyed to engagement of the OMF partner protein during the initial assembly of the tripartite complex, or possibly provide directionality of the L-T-O transition during the efflux cycle, which is compatible with the mechanistic model of RND pump cycling suggested recently (8).

## Materials and methods

### Bacterial strains and growth conditions

All strains were derived from *Salmonella enterica* serovar Typhimurium strain SL1344 (henceforth referred to as *S.* Typhimurium), a pathogenic strain first isolated from an experimentally infected calf (43). All strains were grown in LB broth at 37°C with aeration.

### Growth kinetic assays

Overnight cultures (~10^9^ cfu/mL) of test strains were diluted to a starting inoculum of 10^6^ cfu/mL in a 96-well plate. Where appropriate, the test strains were diluted in LB broth supplemented with antibiotics. Growth was monitored over 12 hr in a FLUOstar OMEGA plate reader (BMG Labtech, Germany).

### Site directed mutagenesis

Mutations in AcrA was generated using the plasmid p*acrA* (pET-20b (+) carrying the *acrA* gene from *S.* Typhimurium SL1334 with C-terminal 6xHis-tag). Mutations in both AcrA and AcrB was generated using the plasmid p*acrAB* (pET-20b (+) carrying the *acrAB* operon from *S.* Typhimurium SL1334 with a C-terminal 6xHis-tag). Mutations in AcrE were generated using the plasmid p*acrE* (pTrcHis2-TOPO carrying the *acrE* gene from *S.* Typhimurium SL1334 with a C-terminal 6xHis-tag). All site-directed mutagenesis (SDM) reactions were carried out using the QuikChange Lightning SDM Kit (Agilent, USA). The mutations were verified by sequencing (Eurofins Genomics, UK). Primers used for all the SDM reactions are listed in Table S1.

### Ethidium bromide accumulation and efflux assay

The efflux activity of strains was assessed by measuring ethidium bromide accumulation and efflux as previously described (44).

### Doxorubicin efflux assay

Doxorubicin efflux was measured in a similar manner to ethidium bromide efflux, with some changes. Cells were grown to an OD_600_ of 0.6 and washed with efflux buffer (20 mM potassium phosphate buffer with 5 mM magnesium chloride) three times. Carbonyl cyanide *m*-chlorophenylhydrazone (CCCP) and doxorubicin were added at a final concentration of 100 μM and 20 μM, respectively. Cells were incubated at 37°C with aeration for 1 hour. Following incubation, cells were washed with efflux buffer three times. Cells were energised with 25 mM glucose and doxorubicin efflux was measured over 30 min at excitation and emission wavelengths of 485 and 620-10 nm, respectively.

### Antimicrobial susceptibility

The agar doubling dilution method was used to determine the minimum inhibitory concentrations (MICs) of various antimicrobials and dyes according to Clinical and Laboratory Standards Institute guidance (45). All MICs were repeated at least three times.

### Western blotting

Wild-type and mutant AcrA were expressed in SL1344 Δ4PAP from p*acrA* plasmids. Wild-type and mutant AcrE were expressed in SL1344 Δ4PAP Δ*acrF* from p*acrE* plasmids. Wild-type and mutant AcrB were expressed in SL1344 Δ4PAP Δ*acrB* from p*acrAB* plasmids. Cultures were grown to an OD_600_ of 0.4 without induction. Cells were harvested and lysed in 10 mM Tris-HCl, 1 mM disodium EDTA, pH 8.0, supplemented with complete EDTA-Free Protease Inhibitor tablets (Roche, Switzerland) and 100 μg/mL lysozyme using sonication. Membrane fractions were harvested, separated using a 12% SDS-PAGE gel for AcrA and AcrE and 8% SDS-PAGE gel for AcrB, and transferred to a PVDF membrane. The His-tagged proteins were blotted using anti-6x His tag HRP-conjugated monoclonal antibody (Invitrogen, USA) and detected using Clarity Western ECL Substrate (Bio-Rad, USA) on an Amersham 680 Imager (Cytiva, USA).

### Molecular visualisation of substrate channels

The location of the substrate channels 1-3 within the RND-transporter AcrB were calculated using CAVER software v3.0 (46) as described previously (26). For visualisation of the recently reported channel 4, we used the CAVER-output kindly provided by K. M. Pos (personal communication), as reported in Tam et al. (28). PyMOL (Molecular Graphics System, Version 2.0 Schrödinger, LLC.) was used for 3D rendering of molecular structures and the substrate channels discussed.

### Statistical analysis

Experiments were carried out at least three times on separate occasions. Data shown are the mean of at least three biological replicates, and where shown, error bars indicate standard deviations. All statistical comparisons were performed using one-way ANOVA with multiple comparisons in GraphPad Prism 9.2 software (GraphPad Software LLC).

## Acknowledgements

I.A. was funded by the Midlands Integrative Biosciences Training Partnership (MIBTP2) and grant BBSRC BB/M01116X/1 at the University of Birmingham. V.N.B. was supported by funding from BBSRC (grant BB/N002776/1). J.M.A.B. was funded by the BBSRC grant BB/M02623X/1 (David Phillips Fellowship to J.M.A.B).

