## Supplementary Material for "The membrane-proximal domain of the periplasmic adapter protein plays a role in vetting substrates utilising channels 1 and 2 of RND efflux transporters"

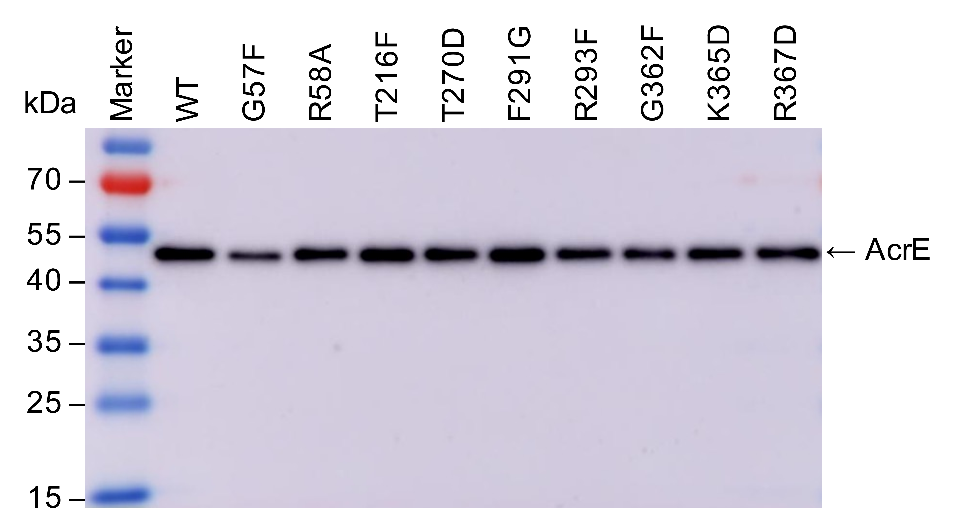

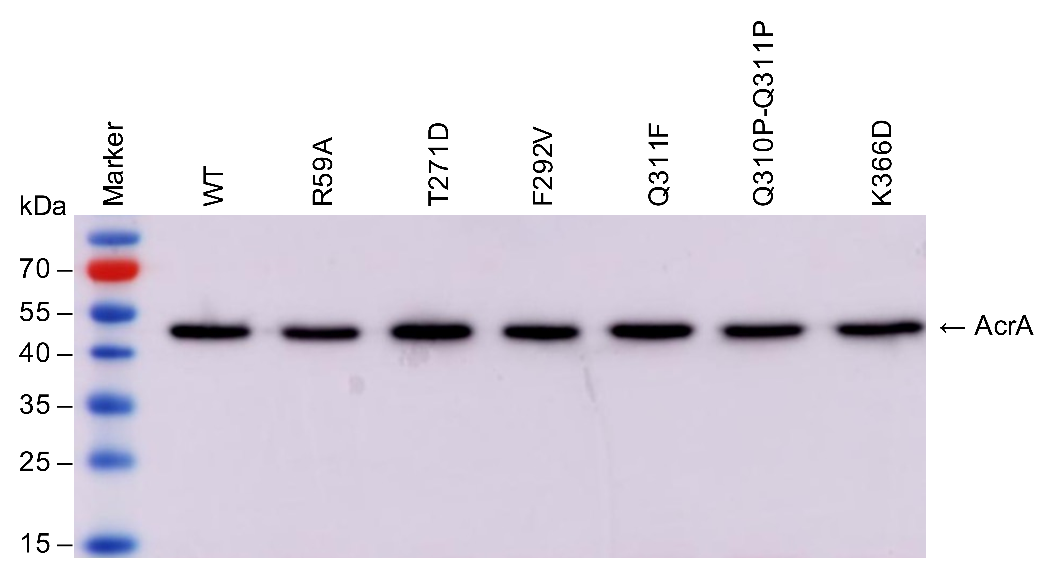

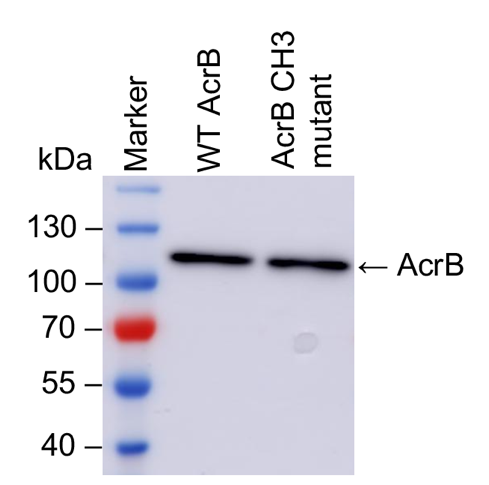

A

B

C

**Figure S1. Western blotting results of mutant proteins. A)** Wild-type and mutant AcrA were in expressed in the SL1344 Δ4PAP strain from p*acrA* plasmids. **(B)** Wild-type and mutant AcrE were in expressed in the SL1344 Δ4PAP Δ*acrF* strain from p*acrE* plasmids. **(C)** Wild-type and the AcrB channel 3 (CH3) mutant were in expressed in the SL1344 Δ4PAP Δ*acrB* strain from p*acrAB* plasmids. Membrane fractions were harvested, separated on a 12% SDS-PAGE gel for AcrA and AcrE or 8% SDS-PAGE gel for AcrB, and transferred to a PVDF membrane. The His-tagged proteins were blotted using anti 6x-His tag HRP-conjugated monoclonal antibody and detected using ECL substrate. PageRuler Prestained Protein Ladder (Thermo Scientific, USA) was used as a molecular weight marker.

**Table S1. List of primers used for site-directed mutagenesis reactions.**

| **Primer name** | **Primer sequence** |
| --- | --- |
| AcrA_R59A_F | CTTCCGGGTGCTACCGTTGCTTACCGTATC |
| AcrA_R59A_R | GCAACGGTAGCACCCGGAAGTTCAGTTGTG |
| AcrA_T270A_F | GTTGACCAAGCCACCGGGTCTATTACTTTG |
| AcrA_T270A_R | CCCGGTGGCTTGGTCAACGGTCACGTCGG |
| AcrA_T270D_F | GACCGTTGACCAAGACACCGGGTCTATTAC |
| AcrA_T270D_R | GACCCGGTGTCTTGGTCAACGGTCACGTCG |
| AcrA_T271A_F | GTTGACCAAACCGCCGGGTCTATTACTTTGC |
| AcrA_T271A_R | GTAATAGACCCGGCGGTTTGGTCAACGGTC |
| AcrA_T271D_F | GTTGACCAAACCGACGGGTCTATTACTTTGC |
| AcrA_T271D_R | GTAATAGACCCGTCGGTTTGGTCAACGGTC |
| AcrA_G272A_S273A_F | CAAACCACCGCGGCTATTACTTTGCGCGCC |
| AcrA_G272A_S273A_R | GTAATAGCCGCGGTGGTTTGGTCAACGGTCAC |
| AcrA_F292V_F | CCAGGAATGGTCGTTCGCGCACGTCTGC |
| AcrA_F292V_R | CGAACGACCATTCCTGGCAATAAGGTGTG |
| AcrA_Q310P_Q311P_F | CTGGTTCCACCACCGGGCGTTACCCGTACTC |
| AcrA_Q310P_Q311P_R | GTAACGCCCGGTGGTGGAACCAGTAATGCCG |
| AcrA_Q311F_F | GTTCCACAATTCGGCGTTACCCGTACTCC |
| AcrA_Q311F_R | GTAACGCCGAATTGTGGAACCAGTAATGC |
| AcrA_P317G_F | GTTACCCGTACTGGACGCGGCGATGCCAC |
| AcrA_P317G_R | TCGCCGCGTCCAGTACGGGTAACGCCCT |
| AcrA_P317F_F | GTTACCCGTACTTTCCGCGGCGATGCCACG |
| AcrA_P317F_R | CGCCGCGGAAAGTACGGGTAACGCCCTGTTG |
| AcrA_R318F_F | CGTACTCCATTCGGCGATGCCACGGTGCTG |
| AcrA_R318F_R | CATCGCCGAATGGAGTACGGGTAACGCCCTG |
| AcrA_I343F_G344F_F | GCCAGGCGTTCTTCGATAAGTGGCTGGTGAC |
| AcrA_I343F_G344F_R | CACTTATCGAAGAACGCCTGGCTTGCGACG |
| AcrA_Q365F_F | GCGGGCTGTTCAAAGTACGTCCTGGCGCAC |
| AcrA_Q365F_R | GACGTACTTTGAACAGCCCGCTGACGACTAC |
| AcrA_K346A_F | AGCCAGGCGATCGGCGATGCGTGGCTGGTG |
| AcrA_K346A_R | CACCAGCCACGCATCGCCGATCGCCTGGCT |
| AcrA_K366D_F | GGCTGCAAGATGTACGTCCTGGCGCACAGG |
| AcrA_K366D_R | GGACGTACATCTTGCAGCCCGCTGACGAC |
| AcrA_R368A_F | CAGCGGGCTGCAAAAAGTAGCTCCTGGCGCA |
| AcrA_R368A_R | TGCGCCAGGAGCTACTTTTTGCAGCCCGCTG |
| AcrA_R368F_F | GCAAAAAGTATTTCCTGGCGCACAGGTTAAAG |
| AcrA_R368F_R | CGCCAGGAAATACTTTTTGCAGCCCGCTGAC |
| AcrE_G57F_F | GTAACGACCGAACTTCCCTTCCGTACGTCCGCATTTCGC |
| AcrE_G57F_R | GCGAAATGCGGACGTACGGAAGGGAAGTTCGGTCGTTAC |
| AcrE_R58A_F | ACCGAACTTCCCGGAGCTACGTCCGCATTTCG |
| AcrE_R58A_R | CGAAATGCGGACGTAGCTCCGGGAAGTTCGGT |
| AcrE_T216F_F | CGATCCGATTTATGTCGACGTGTTCCAATCAAGCAACGACTTTATGC |
| AcrE_T216F_F | GCATAAAGTCGTTGCTTGATTGGAACACGTCGACATAAATCGGATCG |
| AcrE_T270D_F | GTTACCGTAGATGAAAGCGACGGCTCTATCACGCTCAG |
| AcrE_T270D_R | CTGAGCGTGATAGAGCCGTCGCTTTCATCTACGGTAAC |
| AcrE_F291G_F | CTGCTTCCCGGTATGGGTGTTCGCGCCCGCAT |
| AcrE_F291G_R | ATGCGGGCGCGAACACCCATACCGGGAAGCAG |
| AcrE_R293F_F | GTCTGCTTCCCGGTATGTTTGTTTTCGCCCGCATTGA |
| AcrE_R293F_R | TCAATGCGGGCGAAAACAAACATACCGGGAAGCAGAC |
| AcrE_G362F_F | CGATAAGGTCATCGTCAGCTTCTTACAAAAAGCGCGACCG |
| AcrE_G362F_R | CGGTCGCGCTTTTTGTAAGAAGCTGACGATGACCTTATCG |
| AcrE_K365D_F | CATCGTCAGCGGCTTACAAGATGCGCGACCGG |
| AcrE_K365D_R | CCGGTCGCGCATCTTGTAAGCCGCTGACGATG |
| AcrE_R367D_F | CGGCTTACAAAAAGCGGATCCGGGCGTCCAGGTG |
| AcrE_R367D_R | CACCTGGACGCCCGGATCCGCTTTTTGTAAGCCG |
| AcrB_A33W_F | GCGATCCTCAAATTGCCGGTATGGCAATATCCGACGAT |
| AcrB_A33W_R | ATCGTCGGATATTGCCATACCGGCAATTTGAGGATCGC |
| AcrB_T37W_F | GGTATGGCAATATCCGTGGATTGCGCCACCAGCA |
| AcrB_T37W_R | TGCTGGTGGCGCAATCCACGGATATTGCCATACC |
| AcrB_N298W_F | TGGCTACCGGCGCCTGGGCGCTGGATACCGC |
| AcrB_N298W_R | GCGGTATCCAGCGCCCAGGCGCCGGTAGCCA |
| AcrB_R586D_F | GGGCGCAACGCAAGAGGACACGCAAAAAGTGCTG |
| AcrB_R586D_R | CAGCACTTTTTGCGTGTCCTCTTGCGTTGCGCCC |

**Table S2.** Antimicrobial susceptibility of Δ4PAP strain complemented with mutated versions of AcrA.

|  |  |  |  | **MIC (μg/mL)** | | | | | | |
| --- | --- | --- | --- | --- | --- | --- | --- | --- | --- | --- |
| **Strain** | ACR | CLI | CV | DOX | EB | ERY | FA | MB | NOV | R6G |
| WT | 256 | 256 | 64 | 512 | >1024 | 128 | 1024 | >1024 | 512 | >1024 |
| Δ4PAP | 16 | 2 | 2 | 2 | 16 | 2 | 4 | 8 | 1 | 8 |
| WT complement | 64 | 128 | 16 | 64 | 128 | 64 | 256 | 128 | 128 | 128 |
| R59A | **16** | **2** | **2** | **2** | **32** | **2** | **4** | **16** | **2** | **16** |
| T270A | 64 | 128 | 16 | 64 | 128 | 64 | 256 | 128 | 128 | 128 |
| T270D | 64 | 128 | 16 | 64 | 128 | 64 | 256 | 128 | 128 | 128 |
| T271A | 64 | 16 | 8 | 32 | 64 | **8** | **64** | **32** | **8** | **32** |
| T271D | **16** | **2** | **2** | **2** | **16** | **2** | **4** | **8** | **1** | **8** |
| G272A-S273A | 64 | 128 | 16 | 64 | 128 | 64 | 256 | 128 | 128 | 128 |
| F292V | **16** | **2** | **2** | **2** | **16** | **2** | **4** | **8** | **1** | **8** |
| Q310P-Q311P | **16** | **2** | **2** | **2** | **16** | **2** | **4** | **8** | **1** | **8** |
| Q311F | 64 | 128 | 16 | 64 | 64 | 64 | 128 | 128 | 64 | 128 |
| P317G | 64 | 128 | 16 | 64 | 64 | 64 | 128 | 128 | 128 | 128 |
| P317F | 64 | 128 | 16 | 64 | 128 | 64 | 256 | 128 | 128 | 128 |
| R318F | 64 | 128 | 16 | 64 | 128 | 64 | 256 | 128 | 128 | 128 |
| I343F-G344F | 64 | 128 | 16 | 64 | 128 | 64 | 256 | 128 | 128 | 128 |
| Q365F | 64 | 128 | 16 | 64 | 128 | 64 | 256 | 128 | 128 | 64 |
| K366D | 64 | **16** | 16 | **16** | 64 | **16** | **16** | 128 | **16** | 64 |
| R368F | 64 | 128 | 16 | 64 | 128 | 64 | 256 | 128 | 128 | 128 |

Underlined values highlight values for the ΔPAP strain complemented with WT AcrA. Bold values are at least >2-fold different than the parent strain. ACR, acriflavine; CLI, clindamycin; CV, crystal violet; DOX, doxorubicin; EB, ethidium bromide; ERY, erythromycin; FA, fusidic acid; MB, methylene blue; NOV, novobiocin; R6G, rhodamine 6G.

**Table S3.** Antimicrobial susceptibility of Δ4PAP Δ*acrF* strain complemented with mutated versions of AcrE.

|  | **MIC (μg/mL)** | | | | | | | | | | | | |
| --- | --- | --- | --- | --- | --- | --- | --- | --- | --- | --- | --- | --- | --- |
| **Strain** | ACR | BZK | CHL | CLI | CV | DOX | EB | ERY | FA | MB | MIN | NOV | R6G |
| WT | 256 | 64 | 4 | 512 | 64 | 1024 | >1024 | 64 | 1024 | 1024 | 1 | 512 | >1024 |
| Δ4PAP Δ*acrF* | 16 | 4 | 0.5 | 1 | 2 | 2 | 16 | 4 | 4 | 8 | 0.25 | 1 | 8 |
| WT complement | 256 | 64 | 4 | 128 | 32 | 512 | >1024 | 64 | 512 | 1024 | 1 | 512 | >1024 |
| G57F | **16** | **4** | **0.5** | **4** | **1** | **2** | **16** | **4** | **4** | **8** | **0.25** | **2** | **8** |
| R58A | **16** | **4** | **0.5** | **4** | **1** | **2** | **16** | **4** | **4** | **8** | **0.25** | **2** | **8** |
| T216F | 256 | 64 | 4 | 128 | 32 | 512 | 1024 | 64 | 1024 | 1024 | 1 | 512 | 1024 |
| T270D | **16** | **4** | **0.5** | **4** | **1** | **2** | **16** | **4** | **8** | **8** | **0.25** | **1** | **8** |
| F291G | **16** | **4** | **0.5** | **4** | **2** | **2** | **16** | **4** | **8** | **8** | **0.25** | **1** | **8** |
| R293F | **16** | **8** | **0.5** | **8** | **2** | **8** | **64** | **4** | **32** | **64** | **0.25** | **16** | **64** |
| G362F | **32** | **8** | **0.5** | **16** | **2** | **8** | **64** | **4** | **32** | **128** | **0.25** | **16** | **64** |
| K365D | 128 | 64 | 4 | 128 | 16 | 512 | 1024 | 64 | 256 | 1024 | 1 | 256 | 1024 |
| R367D | 256 | 64 | 4 | 128 | 32 | 512 | 1024 | 64 | 512 | 1024 | 1 | 256 | 1024 |

Underlined values highlight values for the ΔPAP strain complemented with wild-type AcrA. Bold values are at least >2-fold different than the parent strain. ACR, acriflavine; BZK, benzalkonium chloride; CHL, chloramphenicol; CLI, clindamycin; CV, crystal violet; DOX, doxorubicin; EB, ethidium bromide; ERY, erythromycin; FA, fusidic acid; MB, methylene blue; MIN, minocycline; NOV, novobiocin; R6G, rhodamine 6G

**Table S4.** Antimicrobial susceptibility of Δ4PAPΔ*acrB* strain complemented with AcrB channel 3 mutant and K366D AcrA.

|  | **MIC (μg/mL)** | | | | | | | | | | |
| --- | --- | --- | --- | --- | --- | --- | --- | --- | --- | --- | --- |
|  | HMMD | | | | PAC | | | | | | |
| **Strain** | ERY | DOX | FA | NOV | | ACR | BZK | CV | EB | MB | R6G |
| WT | 128 | 1024 | 1024 | 512 | | 256 | 64 | 64 | >1024 | 1024 | >1024 |
| Δ4PAP Δ*acrB* | 4 | 2 | 4 | 1 | | 8 | 2 | 2 | 8 | 8 | 8 |
| WT AcrAB | 64 | 64 | 128 | 64 | | 128 | 32 | 16 | 256 | 256 | 128 |
| WT AcrA + AcrB CH3 mutant | 16 | 32 | 32 | 16 | | 32 | 8 | 8 | 32 | 32 | 32 |
| K366D AcrA + WT AcrB | 16 | 16 | 16 | 16 | | 128 | 32 | 16 | 128 | 128 | 64 |
| K366D AcrA + AcrB CH3 mutant | **4** | **2** | **4** | **1** | | **8** | **2** | **2** | **8** | **8** | **8** |

Underlined values highlight values for the ΔPAP Δ*acrB* strain complemented with WT AcrAB. Bold values highlight the MIC values of the K366D AcrA + AcrB CH3 mutant (A33W T37W N298W AcrB) compared to its single mutant parent strains. HMMD, high-molecular-mass drugs; ERY, erythromycin; DOX, doxorubicin; FA, fusidic acid; NOV, novobiocin; PAC, planar aromatic cation; CV, crystal violet; EB, ethidium bromide; MB, methylene blue; R6G, rhodamine 6G; BZK, benzalkonium chloride; ACR, acriflavine

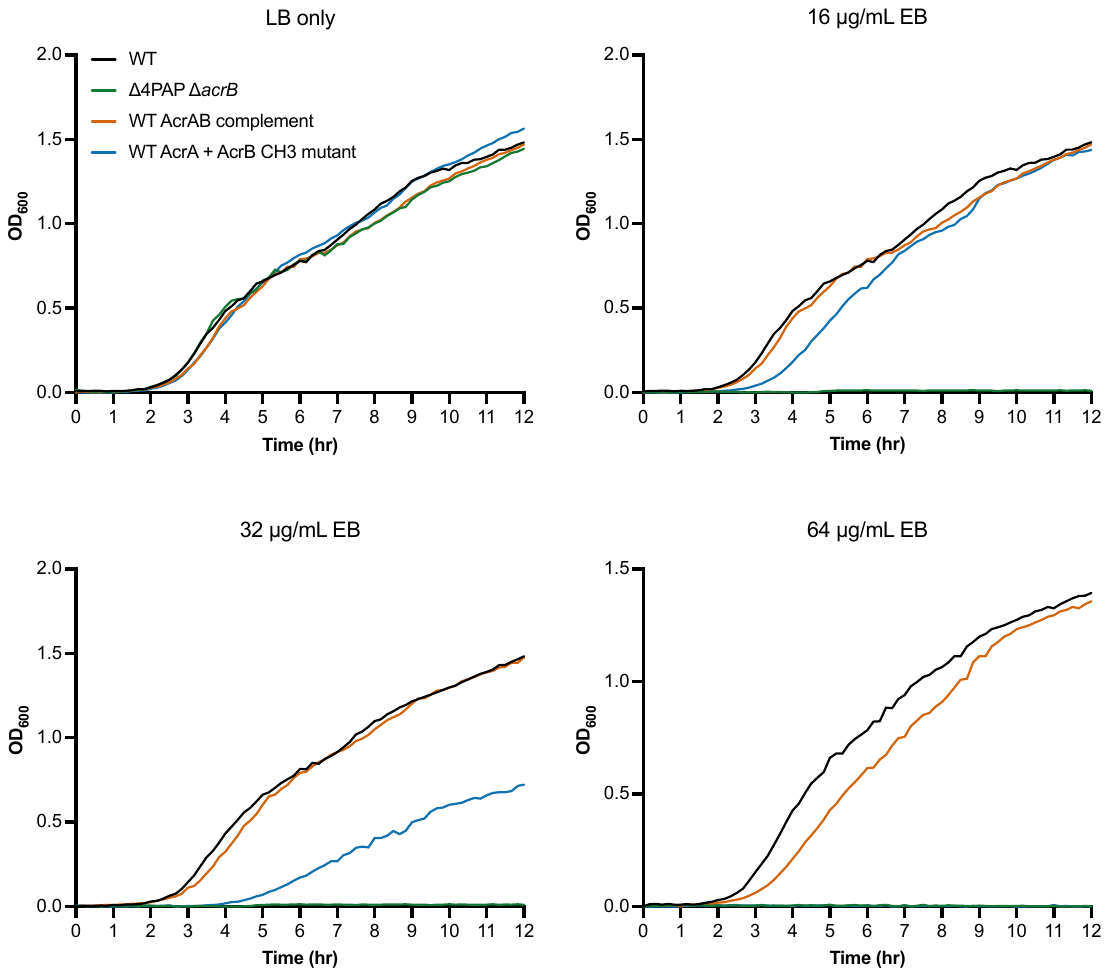

**Figure S2.** Growth kinetics of Δ4PAP Δ*acrB* strain complemented with wild type AcrA and the AcrB channel 3 (CH3) mutant in various concentrations of ethidium bromide (EB). AcrB CH3 mutant refers A33W T37W N298W AcrB. Data shown are the mean OD_600_ values of three biological replicates.

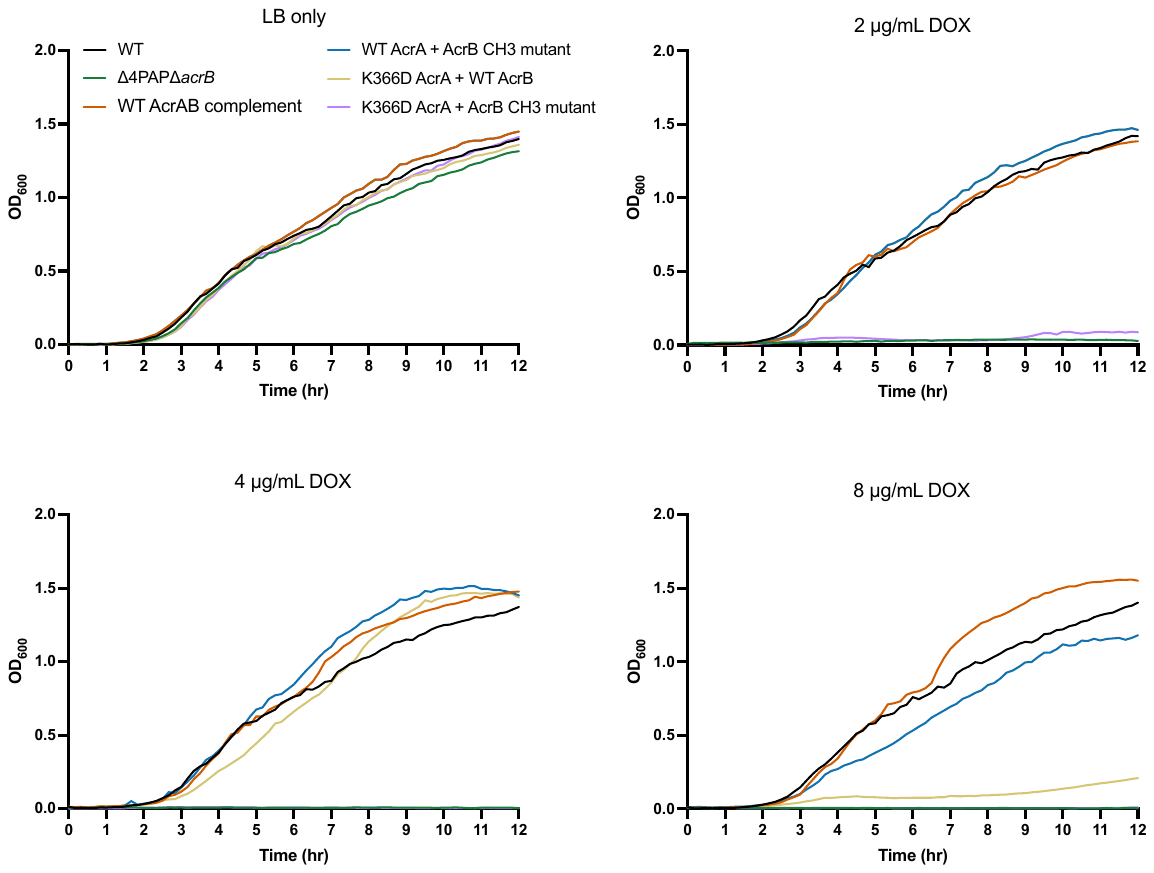
**Figure S3.** Growth kinetics of Δ4PAP Δ*acrB* strain complemented with K366D AcrA and the AcrB channel 3 (CH3) mutant in various concentrations of doxorubicin (DOX). AcrB CH3 mutant refers A33W T37W N298W AcrB. Data shown are the mean OD_600_ values of three biological replicates.

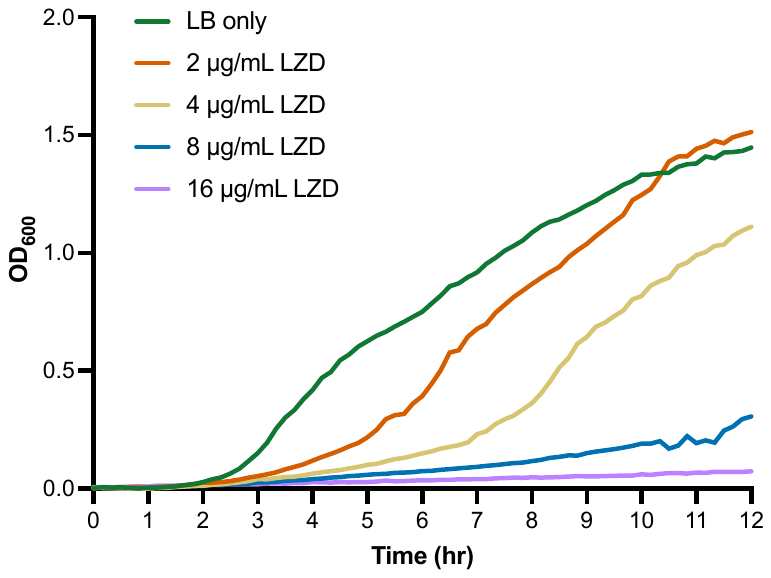

**Figure S4.** Growth kinetics of Δ4PAP strain complemented with K366D AcrA in various concentrations of linezolid (LZD). Data shown are the mean OD_600_ values of three biological replicates.
